# Comparative transcriptomics and host-specific parasite gene expression profiles inform on drivers of proliferative kidney disease

**DOI:** 10.1101/2020.09.28.312801

**Authors:** Marc Faber, Sohye Yoon, Sophie Shaw, Eduardo de Paiva Alves, Bei Wang, Zhitao Qi, Beth Okamura, Hanna Hartikainen, Christopher J. Secombes, Jason W. Holland

## Abstract

The myxozoan parasite, *Tetracapsuloides bryosalmonae* has a two-host life cycle alternating between freshwater bryozoans and salmonid fish. Infected fish can develop Proliferative Kidney Disease (PKD), characterised by a gross lymphoid-driven kidney pathology in wild and farmed salmonids. To facilitate an in-depth understanding of *T. bryosalmonae*-host interactions, we have adopted a two-host parasite transcriptome sequencing approach to minimize host contamination in the absence of a complete *T. bryosalmonae* genome. Parasite contigs common to both infected hosts (the intersect transcriptome; 7,362 contigs) were typically AT-rich (60-75% AT). 5,432 contigs within the intersect were annotated with 1,930 unannotatde contigs encoding for unknown transcripts. We have focused on transcripts encoding proteins involved in; nutrient acquisition, host-parasite interactions, development, and cell-to-cell communication or proteins of unknown function, establishing their potential importance in each host by RT-qPCR. Host-specific expression profiles were evident, particularly in transcripts encoding proteases and proteins involved in lipid metabolism, cell adhesion, and development. We confirm for the first time the presence of homeobox proteins and a frizzled homologue in myxozoan parasites.

The novel insights into myxozoan biology that this study reveals will help to focus research in developing future disease control strategies.

## Introduction

Proliferative Kidney Disease (PKD) is an economically and ecologically important disease that impacts salmonid aquaculture and wild fish populations in Europe and North America^1^. The geographic range of PKD is broad and recent disease outbreaks in a wide range of salmonid hosts underlines the status of PKD as an emerging disease. The occurrence and severity of PKD is temperature driven, with projections of warmer climates predicted to align with escalation of disease outbreaks in the future^2,3^.

PKD is caused by the myxozoan parasite, *Tetracapsuloides bryosalmonae. T. bryosalmonae* spores are released from the definitive bryozoan host, *Fredericella sultana*^4,5^ and, following attachment, invade fish hosts via skin epidermal mucous cells (in gills and elsewhere) and migrate through the vascular system to the kidney and other organs, including spleen and liver^6^. Extrasporogonic proliferation of *T. bryosalmonae* in kidney tissues elicits a chronic tissue pathology, characterised by lymphoid hyperplasia, granulomatous lesions, renal atrophy, anaemia^7,8^ and hyper secretion of immunoglobulins^9,10^. The severity and development of these hallmark symptoms of PKD are modified by a variety of biological, environmental and chemical stressors, impacting on parasite load, host immunity, and disease recovery^10,11^. A genetic basis for such pathological and developmental variation is evidenced by introduced, non-co-evolved salmonid hosts in Europe, such as rainbow trout, acting as dead-end hosts. In contrast to native salmonids, the European strains of *T. bryosalmonae* are unable to produce viable sporogonic renal stages, which are infective to the bryozoan host^5^. Despite these recent advances in understanding the host responses to PKD, the molecular bases of the host-parasite interactions that drive PKD development are currently poorly known and there are no therapeutic measures for disease control.

An increasing number of genomic, transcriptomic and targeted gene studies now show that myxozoans, belong to the Phylum Cnidaria^12–14^. As adaptations to parasitism, myxozoans exhibit extreme morphological simplification and drastically reduced genome sizes relative to free-living cnidarians^15,16^. Nevertheless, polar capsules homologous to the stinging nematocysts of cnidarians have been retained and are used for host attachment. Whilst myxozoans exhibit an apparent streamlining of metabolic and developmental processes compared to free-living cnidarians, not surprisingly, they have retained large numbers of proteases^18^. Likewise, low density lipoprotein receptor class A domain-containing proteins (LDLR-As) are also numerous. This along with the apparent dominance of myxozoan lipases in infected fish, revealed in this and previous studies, suggests that host lipids are an important source of nutrients for myxozoans^18,19^. LDLR-As may, thus, be particularly important in virulence processes in myxozoan parasites, but to date have not been characterised in *T. bryosalmonae. T. bryosalmonae* belongs to the relative species-poor, early diverging myxozoan clade Malacosporea, which has retained primitive features, such as epithelial layers and, in some cases, musculature^20^. Malacosporeans alternate between fish and freshwater bryozoan hosts^17^. There are clear host-specific developmental differences, including meiosis in bryozoan hosts and morphologies of spores released from fish and bryozoan hosts^21^. Host specific differences in gene expression may provide avenues for the development of targeted future therapeutics and are an important prerequisite in understanding the parasite’s biology. However, biological characterization of myxozoans via transcriptome and genome data has for various reasons been hampered. Provision of sufficient and appropriate material may be problematic. For example, sporogonic stages (henceforth referred to as spore sacs) of *T. bryosalmonae* can be released from the body cavity of bryozoan hosts by dissection and can occasionally be collected in substantial quantity (e.g. hundreds of spore sacs) from infected bryozoans maintained in laboratory mesocosms^22^ or from field-collected material^3^. However, attempts to purify parasite stages from infected fish kidney tissues have been unsuccessful. Although dual RNA-Seq approaches and selective enrichment of parasite stages from host tissues can be attempted, most myxozoan genomes and transcriptomes still carry host contamination^23^. *In situ* expression experiments are hindered by low parasite to host tissue ratios in infected tissues and the subsequent very low coverage of parasite transcripts.

In the absence of a complete host-free *T. bryosalmonae* genome, we used transcriptomes from the fish and bryozoan hosts to develop an intersect transcriptome for comparative transcriptomics. After maximising parasite representation in tissue samples of both hosts, normalised cDNA libraries were created to maximise the characterisation of low abundance transcripts. The transcripts expressed in both host transcriptomes were retrieved in the intersect transcriptome and coupled with a novel RTq-PCR assay to assess host-specific expression profiles of parasite transcripts of interest. Our rationale was that different expression profiles are likely to indicate the relative importance of parasite genes and proteins in each host. We particularly focused on proteins linked to putative virulence mechanisms, including; cell/tissue invasion, immune evasion, protein/lipid metabolism, cell-cell communication, and development^15,18,24–27^. We hypothesised that *T. bryosalmonae* uses different virulence strategies in the two hosts and tested this by comparative analysis of the expression profiles of candidate key virulence genes implicated in nutritional, metabolic, virulence and developmental activities. Our approach provides the first transcriptome datasets for *T. bryosalmonae* and affords valuable insights into malacosporean nutritional and metabolic processes and potential virulence candidates. We identify multiple members of key gene families that may be involved in differential exploitation strategies in different hosts. These data have also revealed, for the first time in myxozoans, transcripts encoding homeobox proteins and a frizzled homologue, with the latter implying that a Wnt signalling pathway could be present in myxozoans.

## Results

### Transcriptome assembly and identification of intersect contigs

Sequencing of the normalized cDNA library from bryozoan-derived *T. bryosalmonae* spore sac RNA generated a total of 164,929,000 reads. After quality filtering and adapter trimming, 154,001,594 raw reads were assembled into 48,153 contigs. 87.7% of filtered reads could be re-aligned to the assembly using Bowtie software. Sequencing of *T. bryosalmonae* infected trout kidney RNA generated 160,556,000 reads with 154,576,737 paired reads remaining after sequence trimming. Further sequential filtering was carried out by aligning remaining reads to a draft rainbow trout (*Oncorhynchus mykiss*) genome and a multi-sourced rainbow trout transcriptome database^28^ leaving 57,031,747 and 31,731,181 paired reads respectively. Remaining reads were assembled into 81,035 contigs. 82.9% of filtered reads could be re-aligned to the assembly using Bowtie software.

BLAST analysis of the fish kidney-derived transcriptome assembly against a draft *T. bryosalmonae* genome (Hanna Hartikainen, Beth Okamura, unpublished data) resulted in 5,384 matches (6.6% of the fish kidney-derived transcriptome). Only 191 contigs were filtered with the *Thelohanellus kitauei* genome assembly with all 191 also being filtered with the *T. bryosalmonae* genome. Owing to the low filtering capability of the *T. kitauei* genome assembly, only the draft *T. bryosalmonae* genome was subsequently used to filter parasite contigs from the spore sac-derived transcriptome assembly, yielding 21,720 matches (45.1% of the spore sac-derived transcriptome). Reciprocal BLAST analysis of the myxozoan-filtered spore sac transcriptome against the myxozoan-filtered fish kidney-derived transcriptome yielded 7,362 matches and 14,358 mismatches, whilst the converse revealed 4,723 matches and 661 mismatches. The 7,362 contig matches were defined as the intersect transcriptome. Assembly statistics are shown in Table 1. Initial annotation, based on BLASTX against the nr NCBI database was used to partition each transcriptome dataset into transcripts according to homology with; eukaryotes, bacteria, archaea, and viruses. Remaining contigs with *E* values above the 10^−3^ threshold were deemed as unknown / no hit. In contrast to unfiltered spore sac and fish kidney-derived contig sets, the myxozoan-filtered contig sets were enriched with AT-rich contigs (Fig. 1). The intersect transcriptome is available via figshare (www.figshare.com, 10.6084/m9.figshare.11889672).

**Table 1.**
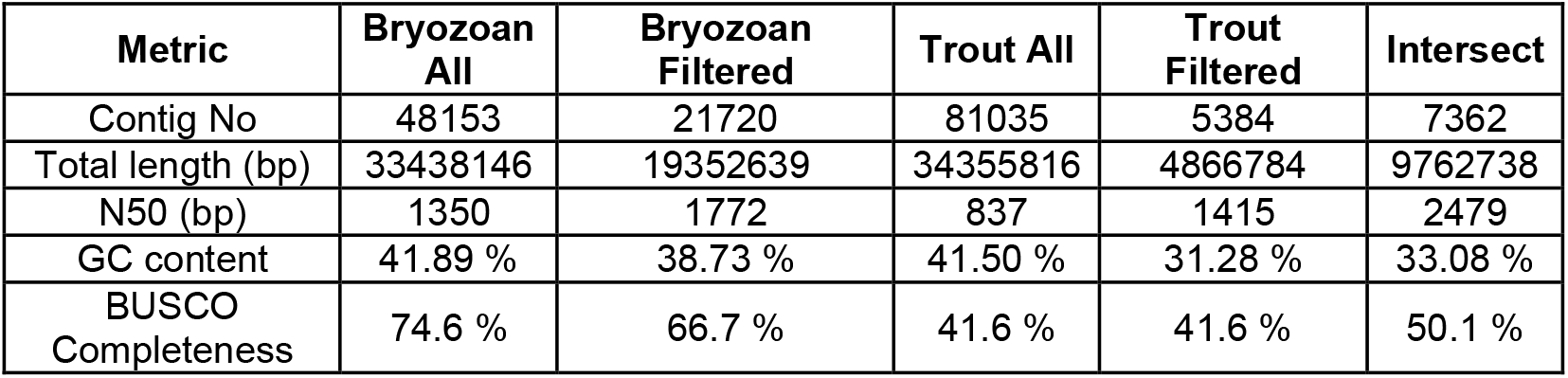
Assembly statistics for sequenced *T. bryosalmonae* transcriptome datasets.

| Metric | Bryozoan All | Bryozoan Filtered | Trout All | Trout Filtered | Intersect |
| --- | --- | --- | --- | --- | --- |
| Contig No | 48153 | 21720 | 81035 | 5384 | 7362 |
| Total length (bp) | 33438146 | 19352639 | 34355816 | 4866784 | 9762738 |
| N50 (bp) | 1350 | 1772 | 837 | 1415 | 2479 |
| GC content | 41.89 % | 38.73 % | 41.50 % | 31.28 % | 33.08 % |
| BUSCO Completeness | 74.6 % | 66.7 % | 41.6 % | 41.6 % | 50.1 % |

**Figure 1.**
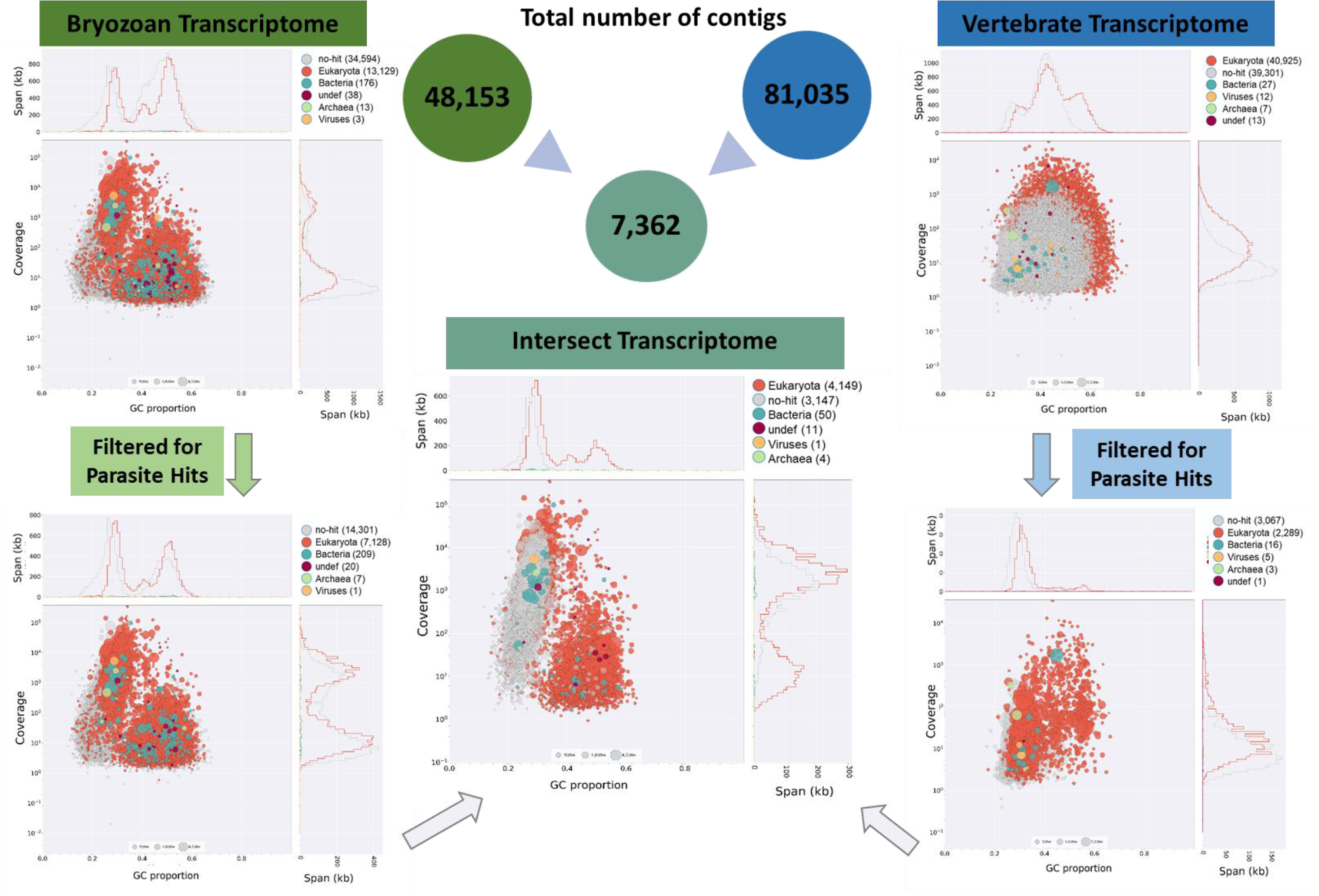
Generation of the *T. bryosalmonae* intersect transcriptome from spore sacs and fish kidney tissues. The contigs derived from each host assembly were filtered against the the *T. kiatauei* genome and/or a draft *T. bryosalmonae* genome to remove host contamination. The intersect transcriptome was generated by BLASTing the filtered spore sac-derived contigs against the fish kidney derived contigs, resulting in 7,362 predicted contigs. The graphs show a BlobPlot of each contig assembly by phylum, with contigs positioned on the X-axis based on their GC proportion and on the Y-axis based on the sum of coverage. Sequence contigs in the assembly are depicted as circles, with diameter scaled proportional to sequence length and coloured by taxonomic annotation.

### Transcriptome annotation, gene ontology and MEROPS analysis

Protein prediction and sequence annotation was performed on the intersect transcriptome as shown in Figure 2. In total, 5432 contigs were homologous to protein sequences in the NCBI database (73.8%) with 1930 (26.2%) deemed as having no homology to known proteins. Of the 5,432 contigs, 58% (3,150 contigs) exhibited high homology (*E* value < 10^−50^) and 42% (2,281 contigs) with matches between *E* values of 10^−5^ and 10^−50^ (Supplementary Table S1). 4,015 were assigned to 9,107 GO terms^29^. Predicted GO annotations were divided into 3 groups based on GO terms: 1) Biological process (3,526), 2) Cellular component (1,962), and 3) Molecular function (3,619). The GO term frequencies were plotted using the Web Gene Ontology Annotation Plot 9 (WEGO, version 2.0)^30^ for the GO-tree level 2, based on their properties and function (Fig. 3) and listed for GO-tree level 2, 3 and 4 (Supplementary Table S2). Of the annotatable and unknown protein sequences, 143 and 84 were predicted to contain a signal peptide respectively (Supplementary Tables S1 and S5).

**Figure 2.**
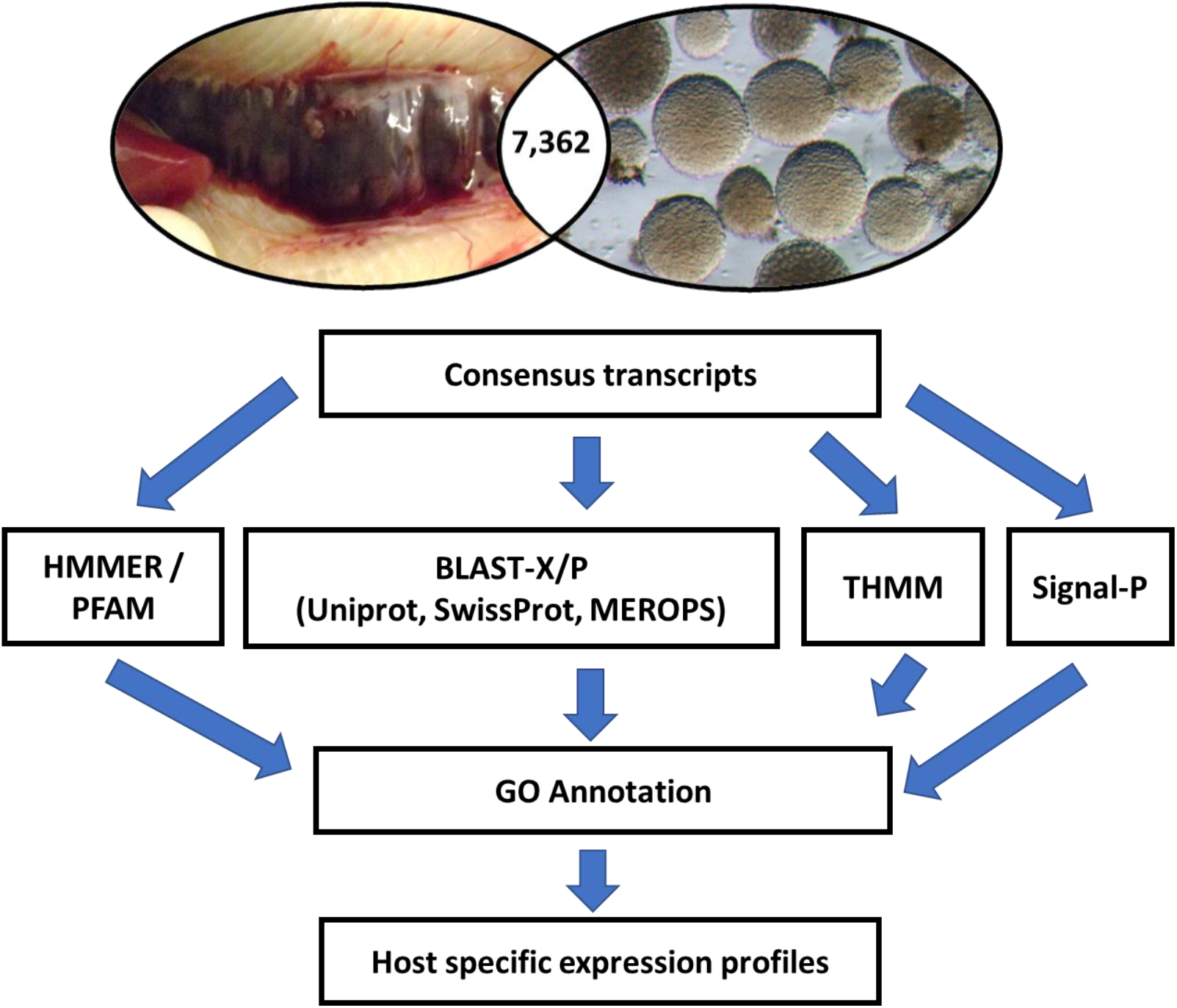
Schematic diagram of the protein annotation process based on the intersect transcriptome from spore sacs and fish kidney tissues. Protein predictions were submitted to the signalP, TMHMM, HMMER and PFam databases and BLAST analysis. Swiss prot matches were mapped to Gene Ontology, EggNOG and KEGG classes. Parasite contigs associated with virulence, metabolism, and development were selected and host-specific expression profiles characterised.

**Figure 3.**
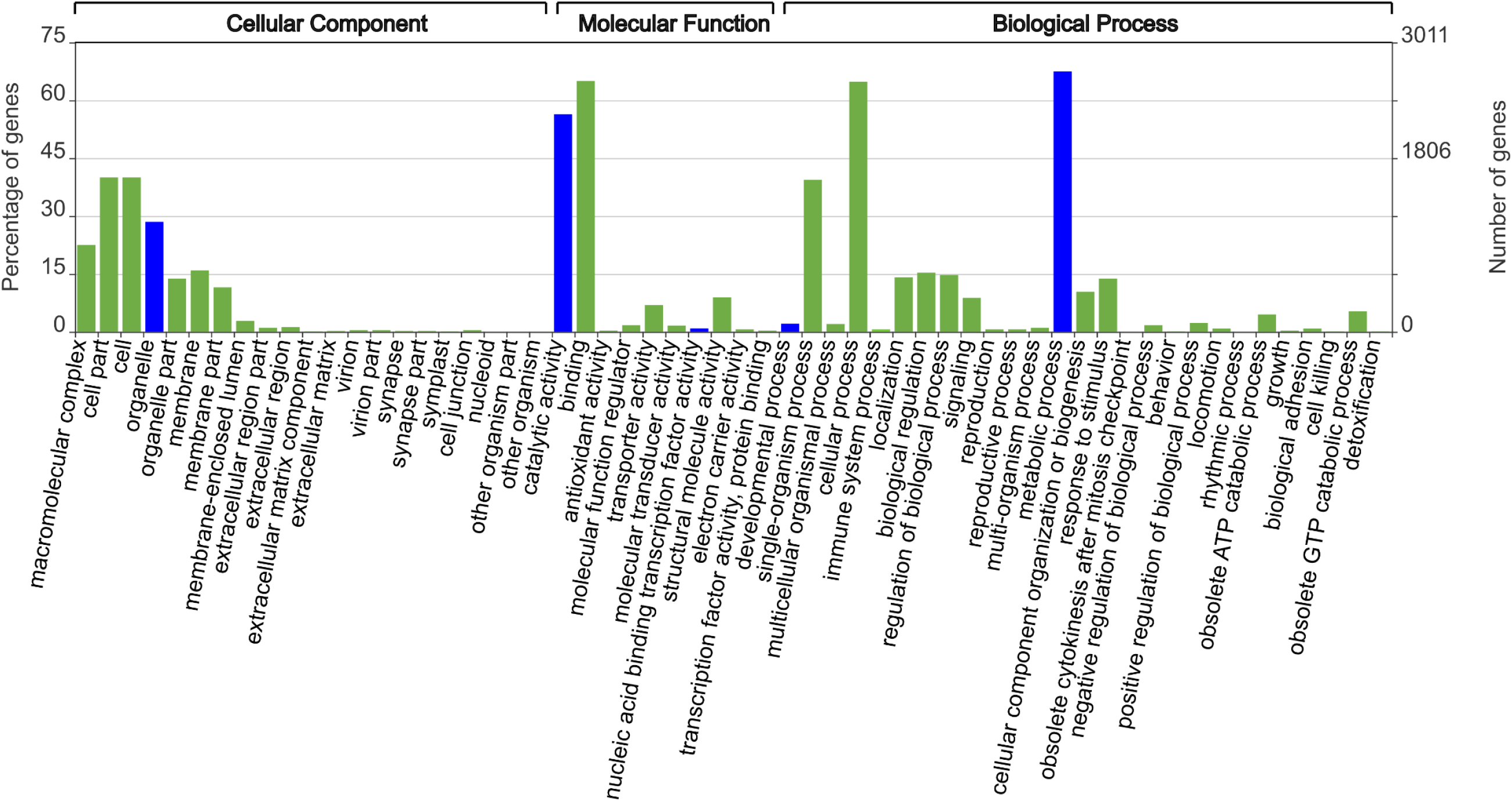
Level 2 Gene Ontology (GO) analysis of *T. bryosalmonae* intersect transcriptome. Functional annotation of the 4,015 annotatable contigs were assigned to 9,107 GO terms and grouped based on their predicted properties and function. The x-axis displays the GO terms selected from the GO trees. The left and right y-axes correspond to percentage gene number and actual gene number per selected GO term respectively. GO terms of relevance to further characterisation in this study are highlighted in blue.

To detect proteases in the intersect transcriptome, all contigs were analysed using the MEROPS database. A total of 100 proteases homologous to metallo-(44) and cysteine (26) proteases, threonine (12), serine (11), aspartate (5) and asparagine proteases (2) were recovered (Supplementary Table S3 and Fig. 4A). All contigs predicted to be lipases or involved in lipid binding based on GO annotation were further manually annotated using BLAST and PFAM. A total of 16 lipases and 22 contigs homologous to lipid binding proteins could be identified (Supplementary Table S4 and Fig. 4B). Of the lipases all, but one, were homologous to phospholipase family members. Specifically, the detected families were phospholipase A1 (8), A2 (4), C (2) and D (1) and a single contig was homologous to the hormone sensitive lipase, lipase E. Two key lipid regulatory enzymes were also uncovered, namely phosphatidate phosphatase and diacylglycerol kinase.

**Figure 4.**
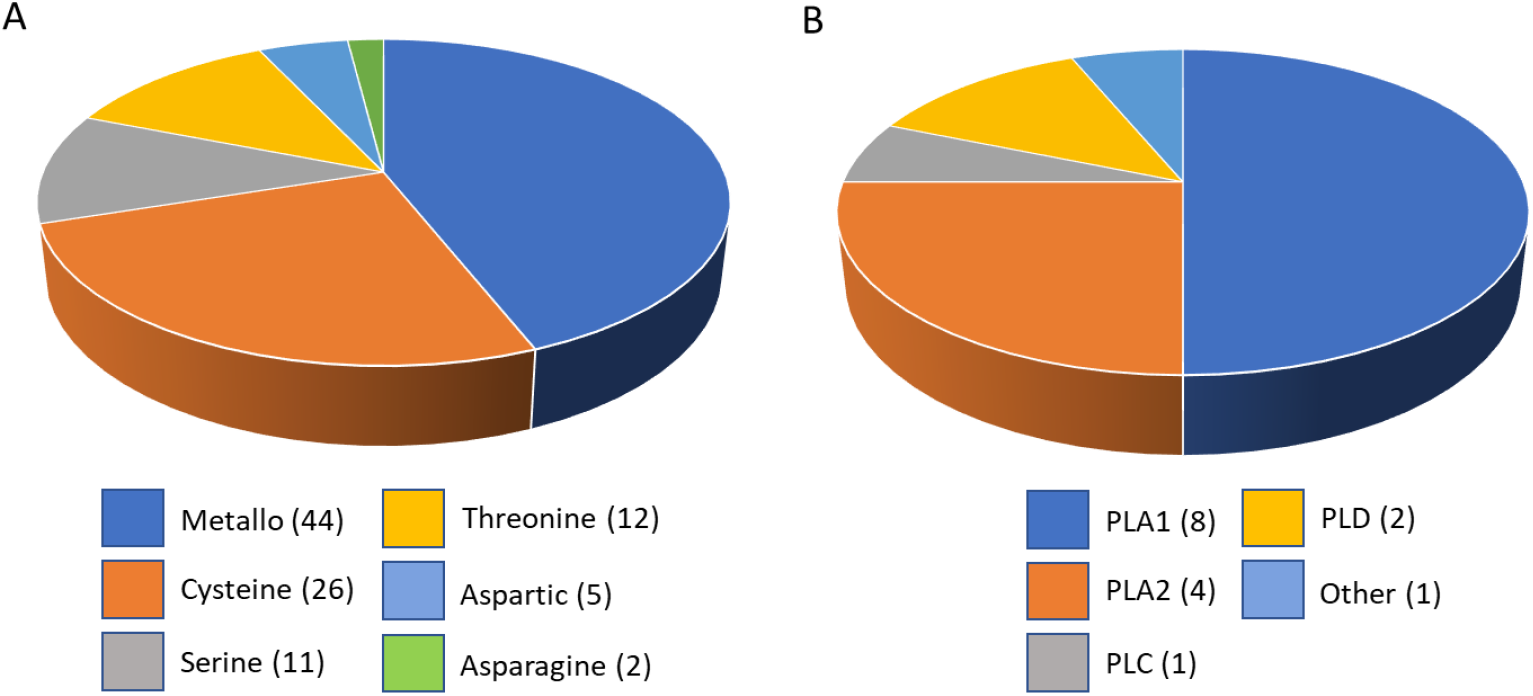
Representation in the intersect transcriptome of A) protease super-families predicted by MEROPS and B) manual annotated and categorised phospholipases (PL). The different colours represent different protease and lipase families.

### Host specific gene expression

The relative expression of 52 selected *T. bryosalmonae* genes were analysed to compare expression profiles in infected bryozoan and infected rainbow trout kidney tissue cDNAs (Figs. 5-8). No expression of any *T. bryosalmonae* gene was evident in control uninfected bryozoan and fish kidney cDNA samples or in -RT controls from uninfected/infected hosts. Three previously known *T. bryosalmonae* house-keeping reference genes were tested, and the mean *Cq* calculated for each host (fish/bryozoan) were: *Tb-*RPL18 (20.00/24.51), *Tb-*RPL12 (20.77/25.03) and *Tb-*ELFα (18.73/23.74). Relative expression normalised to each reference gene individually revealed the same repertoire of significant and non-significant genes. Thus, gene expression data are presented normalised to RPL18, which has been used to determine *T. bryosalmonae* load in trout kidney samples in prevous studies^31^. Δ*Cq* values were calculated representing the expression level of genes of interest compared to *Tb*-RPL18 in each infected host with values >1 indicating lower expression and values <1 indicating higher expression compared to *Tb*-RPL18 (Figs. 5-8). Fold differences in gene expression in infected fish relative to infected bryozoans were assessed for each gene of interest and presented as fold differences in gene expression in Supplementary Table S5. Gene names were assigned according to the closest BLAST search using protein similarity search based on the manually annotated UniProtKB/Swiss-Prot and UniProtKB/Swissprot isoforms databases (https://www.ebi.ac.uk/Tools/sss/fasta/). Highest BLAST hits are listed alongside predicted functional domains (Supplementary Table S5).

**Figure 5.**
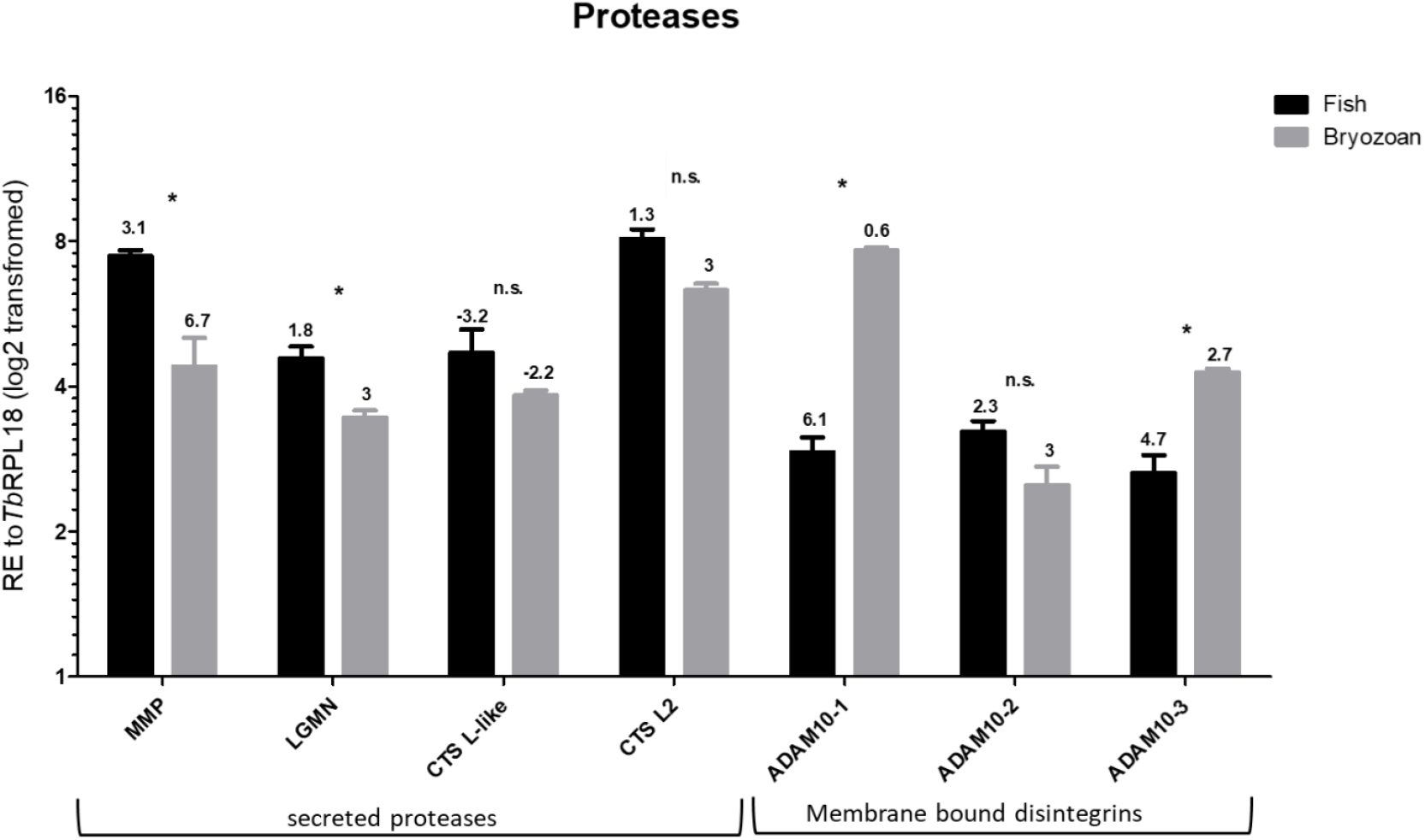
Average relative gene expression in infected bryozoan and infected fish kidney tissues of representative *T. bryosalmonae* proteases. All gene values were normalized to *Tb*-RPL18. Δ*Cq* values, stated above each bar, represent the expression level of genes of interest relative to *Tb*-RPL18 in each infected host with values >1 indicating lower expression and values < 1 indicating higher expression relative to *Tb*-RPL18. \**P*<0.05. Black bars = infected fish, grey bars = infected bryozoan. Gene abbreviations: Matrix metalloprotease 13 (MMP), Legumain (LGMN), Cathepsin L-like (CTLS-like), Cathepsin L2 (CTS L2) and Disintegrin and metalloproteinase domain-containing protein (ADAM) 10-like-1/2/3.

**Figure 6.**
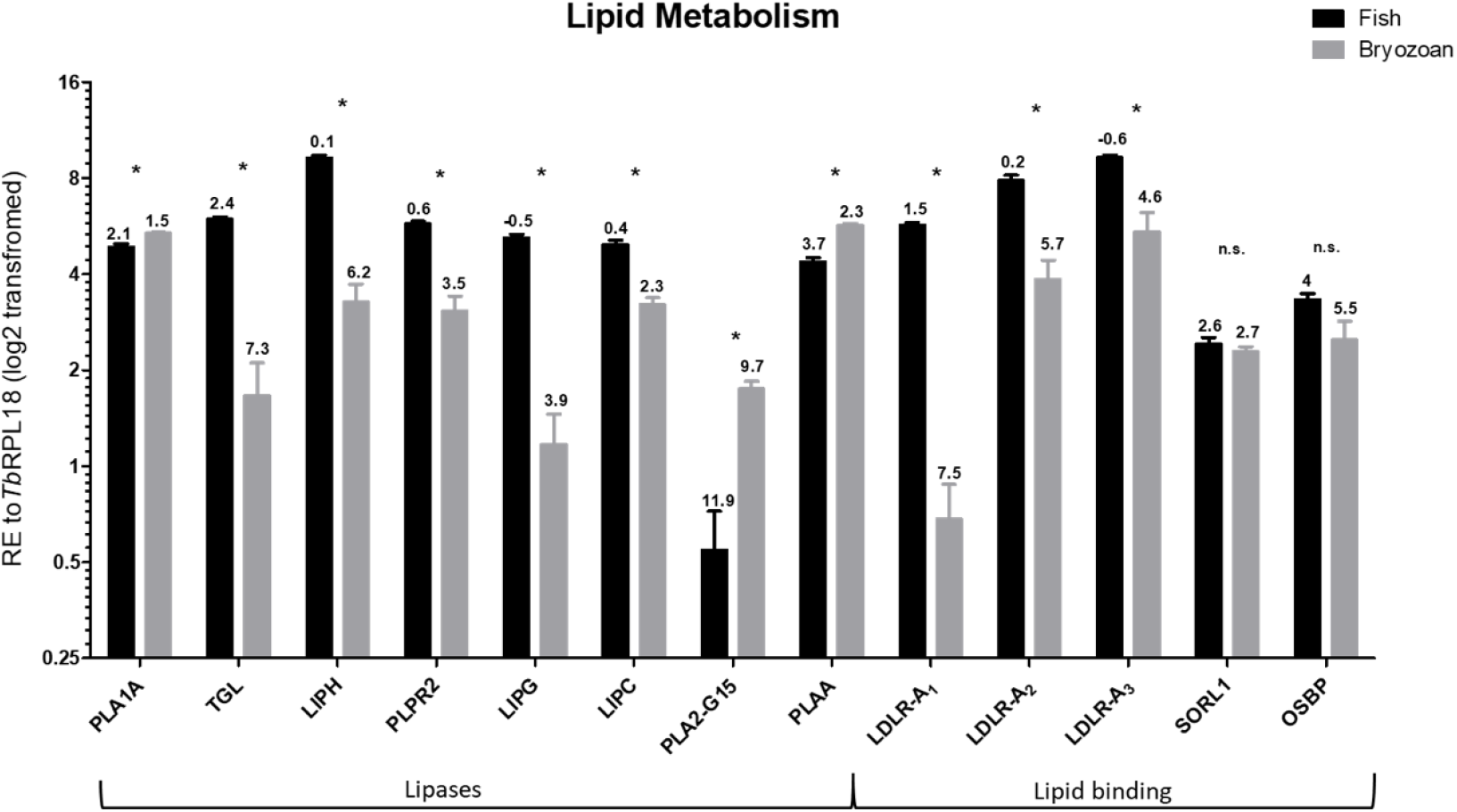
Average relative gene expression in infected bryozoan and infected fish kidney tissues of representative *T. bryosalmonae* genes involved in lipid metabolism. All gene values were normalized to *Tb*-RPL18. Δ*Cq* values, stated above each bar, represent the expression level of genes of interest relative to *Tb*-RPL18 in each infected host with values >1 indicating lower expression and values < 1 indicating higher expression relative to *Tb*-RPL18. \**P*<0.05. Black bars = infected fish, grey bars = infected bryozoan. Gene abbreviations: Phospholipase A1 member A-like (PLA1A), Triacyl glycerol lipase (TGL), Lipase Member H -like (LIPH), Pancreatic lipase-related protein 2 (PLPR2), Endothelial lipase (LIPG), Hepatic triacyl glycerol lipase (LIPC), Group XV phospholipase A2 (PLA2-G15), Phospholipase A-2-activating protein (PLAA), Low-density lipoprotein receptor class A-like protein 1/2/3 (LDLR-A1/2/3), Sortilin-related receptor (SORL1) and Oxysterol-binding protein 2-like (OSBP).

**Figure 7.**
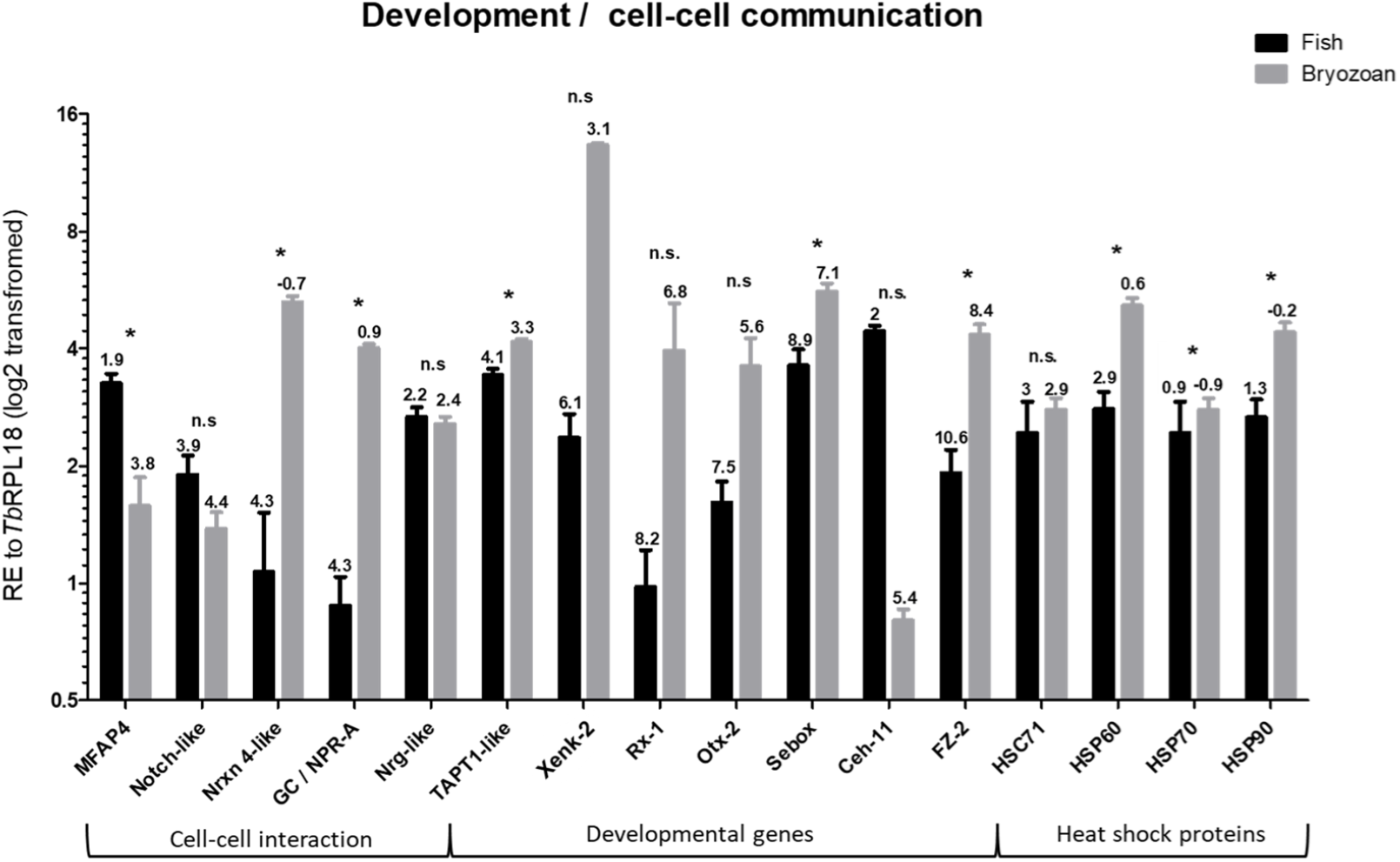
Average relative gene expression in infected bryozoan and infected fish kidney tissues of representative *T. bryosalmonae* genes involved in development and cell-cell communication. All gene values were normalized to *Tb*-RPL18. Δ*Cq* values, stated above each bar, represent the expression level of genes of interest relative to *Tb*-RPL18 in each infected host with values >1 indicating lower expression and values < 1 indicating higher expression relative to *Tb*-RPL18. \**P*<0.05. Black bars = infected fish, grey bars = infected bryozoan. Gene abbreviations: Microfibril-associated glycoprotein 4 (MFAP4), Notch-like protein (Notch-like), Neurexin 4-like (Nrxn 4-like), Guanylate cyclase A / atrial natriuretic peptide receptor A (GC/NPR-A), neuroglian-like protein (Nrg-like), Transmembrane anterior posterior transformation protein 1-like (TAPT1-like), Homeobox protein Xenk-2 (Xenk-2), retinal homeobox-1 (Rx-1), homeobox protein OTX2 (Otx-2), homeobox protein Sebox (Sebox), homeobox protein Ceh-11 (Ceh-11), Frizzled receptor-2 (FZ-2), heat shock protein 60 (HSP60), heat shock cognate 71 kDa protein (HSC71), heat shock protein 70 (HSP70) and heat shock protein 90 (HSP90).

**Figure 8.**
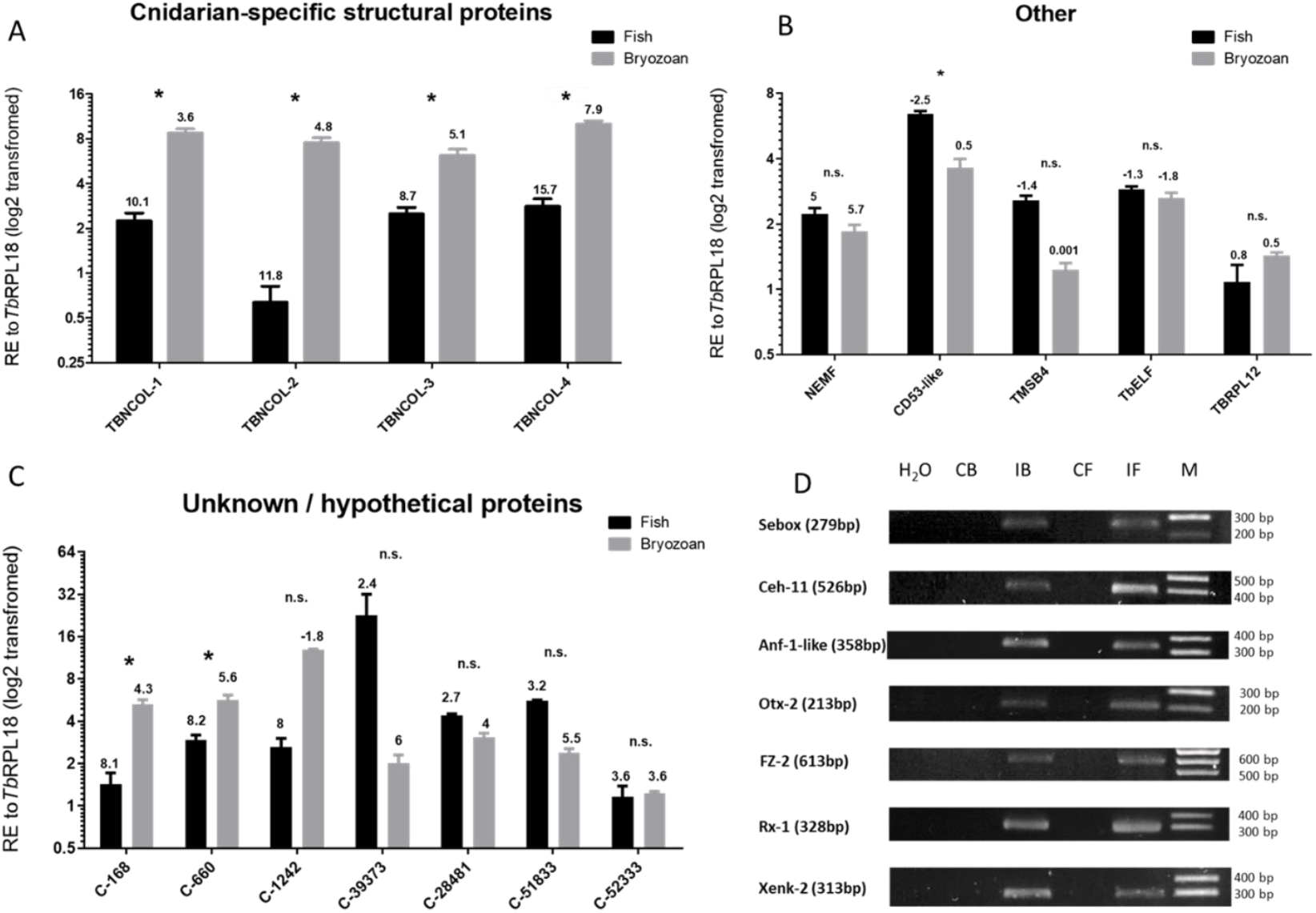
Average relative gene expression in infected bryozoan and infected fish kidney tissues of; A) Cnidarian-specific structural proteins, B) House keeping and immune-related proteins, C) Unknown proteins. All gene values were normalized to *Tb*-RPL18. Δ*Cq* values, stated above each bar, represent the expression level of genes of interest relative to *Tb*-RPL18 in each infected host with values >1 indicating lower expression and values < 1 indicating higher expression relative to *Tb*-RPL18. \**P*<0.05; \*\*\**P*<0.001. Black bars = infected fish, grey bars = infected bryozoan. D) PCR profile showing the presence of Sebox, Ceh-11, Anf-1-like, Otx-2, FZ-2, Rx-1 and Xenk-2 in negative control (H20), uninfected bryozoan and fish kidney controls respectively (CB & CF) and *T. bryosalmonae*-infected bryozoan and fish genomic DNA respectively (IB & IF). Gene abbreviations: Putative group 1 minicollagen-1/2/3/4 (TBNCOL-1/2/3/4), Nuclear export mediator factor (NEMF), Thymosin beta 4-like protein (TMSB4), Tetraspanin CD53-like (CD53-like), *T. bryosalmonae* elongation factor alpha (*Tb*-ELFa) and *T. bryosalmonae* ribosomal protein L12 (*Tb*-RPL12), Unknown proteins C-168, C-660, C-1242, C39373, C-28481, C-51833 and C-52333.

19 genes exhibited significantly higher expression levels in infected bryozoan samples relative to infected fish kidney samples, namely: Disintegrin and metalloproteinase domain-containing protein (ADAM)10-1 (*P*=0.029), ADAM10-3 (*P*=0.029), Phospholipase A1 member A (PLA1A) (*P*=0.029), Phospholipase A-2-activating protein (PLAA) (*P*=0.029), Group XV phospholipase A2 (PLA2-G15) (*P*=0.029), Frizzled receptor-2 (FZ-2) (*P*=0.029), Guanylate cyclase A / atrial natriuretic peptide receptor A (GC/NPR-A) (*P*=0.029), heat shock protein (HSP)60 (*P*=0.029), HSP90 (*P*=0.029), Neurexin 4 (Nrxn4)-like (*P*=0.029), Homeobox protein Sebox (Sebox) (*P*=0.029), Transmembrane anterior posterior transformation protein 1 (TAPT1)-like (*P*=0.029), *T. bryosalmonae* minicollagen-1 (TBNCOL-1) (*P*=0.029), TBNCOL-2 (*P*=0.029), TBNCOL-3 (*P*=0.029), TBNCOL-4 (*P*=0.029), Contig (C)-660 (*P*=0.029), C-168 (*P*=0.029) and HSP70 (*P*=0.029).

In contrast, 12 genes, namely; Matrix metalloprotease 13 (MMP) (*P*=0.004), LDLR-A1 (*P*=0.029), LDLR-A2 (*P*=0.029), LDLR-A3 (*P*=0.029), Legumain (LGMN) (*P*=0.029), Hepatic triacyl glycerol lipase (LIPC) (*P*=0.029), Endothelial lipase (LIPG) (*P*=0.029), Lipase Member H (LIPH) (*P*=0.029), Pancreatic lipase-related protein 2 (PLPR2) (*P*=0.029), Triacyl glycerol lipase (TGL) (*P*=0.029), Microfibril-associated glycoprotein 4 (MFAP4) (*P*=0.029), and Tetraspanin CD53 (CD53)-like (*P*=0.029), exhibited significantly higher expression levels in infected fish kidney samples relative to infected bryozoan samples.

No significantly different expression between the two hosts could be detected for 20 genes, namely; ADAM10-2 (*P*=0.057), Cathepsin L (CTSL)-like (*P*=0.100), CTSL2 (*P*=0.100), Oxysterol-binding protein 2 (OSBP) (*P*=0.114), Sortilin-related receptor (SORL1) (*P*=0.343), Homeobox protein Ceh-11 (Ceh-11) (*P*=0.100), HSC71 (*P*=0.886), Notch-like (*P*=0.114), Neuroglian-like protein (Nrg-like) (*P*=0.486), retinal homeobox-1 (Rx-1) (*P*=0.057), Homeobox protein OTX2 (Otx-2) (*P*=0.057), Homeobox protein Xenk-2 (Xenk-2) (*P*=0.057), Nuclear export mediator factor (NEMF) (*P*=0.200), Thymosin beta 4-like protein (TMSB4) (*P*=0.100), *T. bryosalmonae* elongation factor alpha (*Tb*-ELFa) (*P*=0.200), *T. bryosalmonae* ribosomal protein L12 (*Tb*-RPL12) (*P*=0.343), C-1242 (*P*=0.100), C-39373 (*P*=0.100), C-28481 (*P*=0.100), C-51833 (*P*=0.100) and C-52333 (*P*=0.700).

However, there was a limited availability of biological replicates for infected bryozoan material restricted statistical analysis in some cases (where n=3) and may underly the lack of significant differences (by Mann Whitney U test) for the genes; CTSL-like, Ceh-11, Otx-2, Rx-1, Xenk-2, TMSB4, C-39373, C-1242, C28481, and C51833.

### Presence of frizzled receptor and homeobox genes in *T. bryosalmonae* genomic DNA

To further support the presence of homeobox and FZ-2 genes in *T. bryosalmonae*, primers were designed to specifically amplify genomic DNA from uninfected and infected hosts. PCR analysis detected specific bands, in infected but not in uninfected host tissues, of the correct predicted size and sequence verified for genes; Anf-1-like (358bp), Sebox (279bp), Ceh-11 (526bp), Otx-2 (213bp), FZ-2 (613bp), Rx-1 (328 bp) and Xenk-2 (313bp) (Fig. 8D).

## Discussion

### The intersect transcriptome identifies parasite contigs shared by fish and bryozoan hosts

Comparisons of host-specific expression of key genes may pave the road to targeted approaches to future disease control strategies and identify candidate genes for further functional characterisation. We developed an intersect transcriptome to identify a key set of *T. bryosalmonae* contigs that were expressed to at least some degree in both hosts, whilst minimising host contaminants. In line with our previous studies and existing myxozoan transcriptomes^18,23^, contigs in the intersect transcriptome were predominantly AT-rich (Fig. 1; 60-75% AT). The intersect transcriptome comprised a 7,362 contig group, with a greater number of AT-rich contigs compared to the 5,384 reciprocal contig group but possesses a greater proportion of non-AT rich contigs (Fig. 1). Although low, our Busco scores were comparable to previously reported CEG scores for myxosporeans^15^. It is well known that myxozoan genomes and transcripts are AT-rich (> 60% AT), and AT content can be used to identify transcripts of parasite versus host origin^15,18,23,32^. The increased representation of non-AT-rich contigs in the intersect may, therefore, reflect the presence of non-parasite sequences (see later discussion) but it may also reflect the presence of myxozoan transcripts that encode proteins rich in amino acids using GC rich codons, such as minicollagens^33^. Whilst parasite origin of certain non-AT rich cnidarian/myxozoan-specific transcripts may be identified, others would need to be PCR verified in the absence of a complete host-free parasite genome.

### Host-specific transcription indicates specialisation in nutrient acquisition and virulence

To identify host specific expression patterns, a subset of 52 contigs of specific interest to host-parasite interactions were chosen for RT-qPCR analysis in multiple infected fish and bryozoan samples. Primers designed to these contigs successfully amplified in both infected hosts and in multiple isolates and not in uninfected controls, confirming their parasite origin, along with their AT-rich composition. Proteases have been extensively targeted for therapeutic intervention^25,34,35^ due to their central roles in virulence mechanisms, including nutrient acquisition. Parasite cysteine proteases were represented in the intersect transcriptome (Supplementary Tables S1 and S3). Host protein degrading CTS proteases are often co-expressed with nutrient acquiring and protease activating LGMN-like cysteine proteases in parasites^36^. Similar legumain and CTSL expression profiles and high expression levels in both the fish and bryozoan hosts observed in this study imply that they are biologically important in both infected hosts (Fig. 5).

The dominance of metalloproteases in our *T. bryosalmonae* dataset is not surprising as modulation of the fish kidney tissue matrix, including parasite migration, proliferation and tissue remodelling, are prominent features of PKD^37^. ADAM family members that are active proteases are classified as ectodomain sheddases that regulate signalling associated with extracellular matrix homeostasis in association with tetraspanins. They therefore regulate a wide range of functions including; cell adhesion, inflammation, lymphocyte development, tissue invasion, and notch receptor processing. Consequently, dysregulation of ADAMs is strongly associated with infection-mediated tissue pathology and cancer, implicating both ADAMs and tetraspanins as important therapeutic targets^35,38–40^. In this study, we uncovered six ADAM-like transcripts, with three that are ADAM 10-like and exhibiting variable host expression profiles (Supplementary Table S3 and Fig. 5).

We have also uncovered several tetraspanins with a particularly abundant CD53-like transcript exhibiting a fish-specific expression profile (Supplementary Table S1 and Fig. 8B). In mammals, tetraspanins in the CD family are important immune regulators with hematopoietic-specific CD53 being a suppressor of inflammatory cytokines^41,42^. Thus, association between *T. bryosalmonae* ADAM and tetraspanin homologues may, in part, be responsible for *T. bryosalmonae-*mediated disease pathology and dampening of fish pro-inflammatory responses observed during PKD^11,43^. Similarly, the T lymphocyte hormone TMSB4, dampens inflammatory processes by influencing TLR activity. The very high expression levels of a *T. bryosalmonae* TMSB4 homologue in each host exceeding RPL18, highlight its general biological importance in both hosts.

Parasites often depend on salvaging of host lipids to fuel growth and survival in a range of hosts, including fish^44–46^. It is, therefore, not surprising to find a repertoire of *T. bryosalmonae* phospholipases with the majority belonging to the PLA1 family (Supplementary Table S4). PLA1s hydrolyze acylglycerols and phospholipids, liberating free fatty acids and lysophospholipids^47^. All full-length PLA1 transcripts revealed in this study are likely to be secretory with most having highly fish-specific expression profiles. Importantly, PLA1A and Lipase H release the lipid mediators lysophosphatidylserine (LPS) and lysophosphatidic acid (LPA) that induce cell proliferation and chronic disease pathology^47,48^. It is, therefore, tantalizing to speculate that such lipid mediators may drive PKD pathology. PLA2 activity, can also lead to the release of these lipid mediators, albeit to a lesser extent than PLA1. We did not uncover any calcium dependent PLA2 family members that are linked to arachidonic acid release and production of inflammatory eicosanoids. All PLA2s we found were homologous to intracellular calcium independent PLA2s implicated, to a lesser extent, in immune regulation^49^. Interestingly, PLA2-G15 and PLAA were more highly expressed in bryozoans than in fish kidney tissue suggesting that these family members play a less important role in lipid metabolism of the parasite when in fish than in bryozoan hosts.

As in previous myxozoan studies, we uncovered several LDLR-A transcripts^18^ with 3 being highly fish-specific (Supplementary Table S4 and Fig. 6). LDLR-A proteins are crucial in receptor-mediated endocytosis of ligands, particularly cholesterol and acyglycerols. Many parasites are unable to produce cholesterol and are, therefore, dependent on host acquisition to fuel cell proliferation and differentiation^46,50^. Consequent alterations in host cholesterol homeostasis can lead to immune dysregulation and disease pathology. The apparent dominance of myxozoan LDLR-As and PLA1 family members in infected fish kidney tissue is strongly indicative of host cholesterol and acyglycerol exploitation that could be a major contributory factor to PKD pathogenesis. Indeed, recent studies describe several viable pharmacological approaches to interfere with parasite utilisation of host lipids, which may have relevance to the future control of myxozoan parasites^51–53^.

### Host-specific transcription linked to parasite developmental processes

To develop a fuller understanding of myxozoan biology and potential future directions in disease mitigation, we examined datasets for genes involved in parasite development and functional morphology in each host. ADAM family members, especially ADAM 10, are crucial in the regulation of notch signalling, implicating the involvement of some members in regulation of embryonic and adult developmental processes and thus as potentially important therapeutic targets^38,54^. The presence of meiosis in *T. bryosalmonae* stages in bryozoan hosts, may explain the bryozoan dominance of two of the ADAM 10-like molecules in this study.

Activation of natriuretic peptide receptors, including GC/NPR-A, leads to the accumulation of intracellular cGMP. They act as environmental sensors involved in the regulation of developmental events, blood pressure regulation, immune and electrolyte homeostasis^55^. Cyclic nucleotide balance is also crucial for parasite development, host-to-host transmission, target tissue recognition, and immune evasion, enabling rapid adaptation to different host environments^56,57^. The bryozoan-specific nature of GC/NPR-A in this study may imply similar roles in regulating developmental events leading to transmission and invasion of *T. bryosalmonae* from bryozoans to fish.

Heat Shock Proteins (HSPs) have also been linked to both virulence and parasite development^58^. Significantly higher expression of Hsp60 and Hsp90 in bryozoans in this study may indicate a more important role for these HSPs in infected bryozoans relative to infected fish. Transcripts involved in cell-cell communication such as; neural sensory processing (Nrxn 4-like) and extracellular matrix interactions (MFAP4) exhibited host-specific expression profiles (Fig. 7). The expression of genes linked to neural development is intriguing given the lack of defined nervous systems in myxozoans^20^ and the inert nature of *T. bryosalmonae* sacs.

Minicollagens are likely to play a similar role to those in free-living cnidarians providing the high tensile strength needed to facilitate polar filament deployment^59^. In myxozoans this enables spores to attach to hosts. The dominance of minicollagen expression in bryozoan hosts that support spore development relative to the dead-end fish host is, therefore, an expected result confirmed here for the first time by our expression analyses.

For the first time in myxozoans, we have detected several homeobox proteins and a frizzled receptor homologue (FZD-2). As in previous myxozoan studies, we did not find any hox genes nor other elements of Wnt pathways. *T. bryosalmonae* FZD-2 and most homeobox proteins displayed bryozoan-specific expression profiles. This may further illustrate a developmental deficit in rainbow trout hosts (in contrast to natural fish hosts, such as brown trout, where viable spore development occurs). It is also possible that multicellular sac development in bryozoans requires more sophisticated regulation than cellular proliferation and development of simple pseudoplasmodia in fish hosts. Nevertheless, fish-specific expression of a Ceh11-like transcript suggests that some *T. bryosalmonae* homeobox proteins may have distinct roles in driving fish-specific development. We note, however, that caution must be exercised in assignment of *T. bryosalmonae* homeobox proteins to specific families. This requires in-depth phylogenetic analysis and will be facilitated when more family members are discovered in other myxozoans. Likewise, whether our frizzled receptor is truly indicative of Wnt pathways in myxozoans awaits clarification, particularly as they are also involved in non-Wnt signalling pathways^60^.

### Limitations and caveats

Limitations of this study include lack of insight on proteins expressed during normal development of *T. bryosalmonae* pseudoplasmodia in kidney tubules of natural fish hosts and in the cryptic stages that develop in association with the body walls of bryozoan hosts. A range of other proteins may be expressed during these phases of the life cycle of *T. bryosalmonae*.

Non-parasite (< 60% AT) sequences were evident in the blob plots of both the fish kidney-derived and spore sac-derived transcriptomes. As highlighted by Foox and colleagues^23^, this illustrates the inadequacy of a single step depletion of host sequences by BLAST analysis against fish sequence resources. Notably the second peak in the blob plot of the spore sac-derived transcriptome was not substantially reduced following filtering using a partial *T. bryosalmonae* genome assembly (H. Hartikainen pers comm). This is not surprising as both the genomic and transcriptomic contigs were derived from sequencing libraries prepared from parasite spore sacs isolated from bryozoans and likely carried some bryozoan tissue. Indeed, It is likely that non-parasite sequences are universally present in existing myxosporean assemblies due to the presence of host cell/tissue contaminants^23^. We propose that in the absence of clean and complete host and parasite genomes to filter against, the intersect transcriptome approach can identify a key set of contigs for targeted analysis as shown in this study.

The intersect transcriptome is, however, not assumed to represent a complete transcriptome. A perhaps more important limitation is that numerous parasite transcripts are likely to be amongst the apparent bryozoan-specific and fish-specific mismatches. Both sets of mismatched contigs are unlikely to reflect host exclusivity but rather low transcriptome coverage, limitations of *de novo* assembly, and limited availability of *T. bryosalmonae* resources. Yet, this can also be construed an advantage when comparative follow up studies are conducted, as tracking transcripts expressed in both hosts could help to unravel host-specific aspects of the parasite’s biology. Indeed, BLASTX annotation of both groups of mismatches enabled us to pinpoint potentially important *T. bryosalmonae* players in host-parasite interactions (eg. TMSB4, MMP, LGMN, ADAM10-3; Supplementary Table S5). In addition, all *T. bryosalmonae* homeobox proteins reported in this study were identified in the two groups of host-specific mismatches. Likewise, FZ-2 was identified in the bryozoan-specific group of mismatches (Table 2, Supplementary Table S5). Given the contentious nature of the existence of homeobox and Wnt signalling components in myxozoans, we additionally further confirmed parasite origin via the detection of each gene specifically in genomic DNA from infected hosts (Fig. 8D).

**Table 2.** Predicted developmental genes in *T. bryosalmonae* with closest manually annotated sequence by species given alongside predicted protein domains, namely; signal peptide (SP), frizzled (Fz) and smoothened (Sm).

| Accession Number | Gene name (contig number) | Closest species (Accession No) | E value (BlastX) | % A T | ORF (AA) | Protein domains / motifs |
| --- | --- | --- | --- | --- | --- | --- |
| MN056840 | Homeobox protein, Xenk-2 (C-2190) | Amphibalanus Amphitrite (KAF0307692.1) | 4e-17 | 75 | 196 (FL) | 1x Homeobox |
| MN056845 | Homeobox protein, Anf-1-like (C-8582) | Gallus gallus (P79775) | 5e-12 | 73 | 199 (FL) | 1x Homeobox |
| MN056841 | retinal homeobox-1 (Rx-1) (C-972) | Mus musculus (P80206) | 9e-08 | 72 | 182 (FL) | 1x Homeobox |
| MN056842 | Homeobox protein, OTX-2 (C-18561) | Mus musculus (P80206) | 1e-13 | 68 | 345 (FL) | 1x Homeobox |
| MN056843 | Homeobox protein, Sebox (C-4608) | Callorhinchus milii (XP_007894674) | 6e-02 | 73 | 213 (FL) | 1x Homeobox |
| MN056844 | Homeobox protein, Ceh11 (C-44466)* | Caenorhabditis elegans (P17486) | 1e-12 | 71 | 93 (P) | 1x Homeobox |
| MN056859 | Frizzled receptor-2 (C-2600) | Caenorhabditis elegans (G5ECQ2) | 2e-17 | 72 | 613 (P) | Sp, 1x Fz domain, 1x Fz/Sm membrane region |

Some 26% of contigs could not be annotated and thus encode unknown or orphan proteins. Comparable proportions of unknowns have been reported in other parasite transcriptomic datasets, including myxosporeans^18,23,61^. Myxozoans have have undergone both extreme genome reduction and rapid molecular evolution and the high proportions of unknown proteins may reflect long branches that preclude resolving relationships to currently characterised proteins. Of the seven unknowns examined in this study, five exhibited host-specific expression profiles. It will be a major opportunity and challenge to determine how these unknown proteins may reflect adaptations to parasitism in myxozoans.

### Concluding remarks

We have generated the first insights on malacosporean gene expression based on the analyses of transcriptome datasets using a 2-host sequencing approach. Subsequent RTqPCR analysis enabled us to gain novel insights on myxozoan biology, particularly regarding virulence, metabolism and development. This study will promote future work to understand and mitigate myxozoan-borne diseases.

## Materials and Methods

### Bryozoan and rainbow trout kidney tissue sampling

Infected bryozoans were collected in late spring from the River Cerne, Dorset, UK as described previously^3^. Immature and mature parasites (spore sacs) were isolated by dissection from hosts, rinsed in fresh river water, taken up by capillary action and placed directly into ice-cold Trizol-reagent (Sigma). 150 spore sacs were pooled and used to generate sufficient total RNA for cDNA library construction. Lysed spore sacs in Trizol-reagent were stored at −80 °C prior to RNA extraction.

For gene expression analysis, uninfected and overtly infected bryozoans from the Furtbach River, Switzerland, were used. Bryozoan collections were cultured in a laboratory mesocosm, as described previously^62^, with successfully attaching bryozoan branches constituting a single colony. All bryozoan cultures were maintained for 1-3 weeks to allow colonies to develop transparency, enabling the detection of overt infections using a stereomicroscope. Uninfected bryozoan colonies were assessed based on colony morphology, absence of overt *T. bryosalmonae* spore sacs and by 18S rDNA-based PCR analysis, as described previously^63^. Similarly, confirmation that each spore sac-containing bryozoan sample was, indeed, infected with *T. bryosalmonae*, was obtained by 18S rDNA and RPL18 PCR and sequence verification, as described previously^31,63^. We were unable to distinguish between different stages of overt spore sac infection in bryozoans with only bryozoan colonies exhibiting clear spore sac infection used for gene expression analysis. To ensure sufficient total RNA for RT-qPCR analysis, 4-8 individual uninfected or overtly infected colonies were pooled to form a single biological replicate. In total, 3-4 biological replicates were created for each infection category, with each replicate placed into 1 ml RNAlater for 24 h at 4 °C and stored at −80 °C prior to RNA extraction.

Uninfected and infected rainbow trout exhibiting advanced clinical disease (kidney swelling grade 2 to 3), as determined using the Clifton-Hadley grading system^64^, were sampled from a commercial trout farm located in Southern England with a history of annual PKD outbreaks, as described previously^31^. From parallel studies in our laboratory, examining the dynamics of *T. bryosalmonae* transcript expression at different stages of clinical disease, expression levels in fish kidney samples reached maximal thresholds at grades 2 and 3. Thus, only fish kidneys graded from 2 to 3 were considered in subsequent cDNA library preparation and gene expression analysis (M. Faber, unpublished data). Presence/absence of *T. bryosalmonae* was confirmed by 18S rDNA/ RPL18 PCR and sequence analysis, with extrasporogonic parasite stages in kidney interstitial tissue confirmed by histological examination, as described previously^31,63^. Approximately 100 mg trunk kidney tissue was removed immediately below the dorsal fin from each fish and archived in 1 ml RNAlater, as outlined above.

### RNA extraction and sequencing

Total RNA from *T. bryosalmonae* spore sacs and infected kidney tissue in Trizol-reagent was extracted from Trizol-reagent according to the manufacturer’s instructions. Owing to the presence of PCR inhibitors in bryozoan tissue, bryozoan total RNA was treated with a PowerClean Pro RNA Clean-up kit (Cambio) to remove free and nucleic acid-bound PCR inhibiting substances. All total RNA samples were also treated with Dnase-I using a TURBO DNA-free kit (Invitrogen), quantified using NanoDrop^®^ ND-1000 UV-vis spectrophotometer and quality checked using an Agilent Bioanalyser 2100. cDNA library construction, RNA sequencing, filtering of fish-derived reads using rainbow trout genome and transcriptome databases, and *de novo* assembly was performed by Vertis Biotechnologie AG (Friesing, Germany). Poly (A) + RNA was isolated from 3 µg of spore sac total RNA, transcribed into cDNA, amplified using the Ovation RNA-Seq System V2 kit (NuGEN Technologies Inc.) according to the manufacturer’s instructions and fragmented by sonication. After end-repair and ligation of TruSeq adapters, cDNA was amplified by PCR and normalized by controlled denature-re-association to ensure maximum *T. bryosalmonae* representation in the transcriptome assembly. Single-stranded cDNA was purified by hydroxylapatite chromatography by PCR amplification. Normalized cDNA was gel fractionated and fragments ranging from 300-500 bp in length were sequenced using an Illumina HiSeq 2000 machine producing paired end 100 bp read lengths.

To ensure maximum *T. bryosalmonae* representation in the final fish kidney-derived transcriptome, total RNA from 65 kidney tissue samples was isolated and subjected to RT-qPCR, as described previously^65^. Primers unique to *T. bryosalmonae* and rainbow trout RPL-18 (Supplementary Table S5) were used to determine the most heavily infected kidney samples for cDNA library construction, as described previously^31^. ΔCq values varied from 17.21 (lowest parasite burden) to 5.19 (highest parasite burden). Equal amounts of total RNA were combined from 3 kidney samples with the lowest ΔCq values (5.19, 5.22, 5.95), then cDNA library construction and poly (A) + RNA sequencing performed, as described above.

### Transcriptome Assembly

Raw sequencing reads were quality filtered using a phred quality score cut off-of 33 in ASCII format in accordance with Sanger FASTQ format and a minimum length cut-off of 20 bp using trimming and sequence quality control tools within the CLC Bio Genomics Workbench (version 6.5.1). In the absence of a high-quality *T. bryosalmonae* genome scaffold, sequence reads were assembled *de novo*. In the case of the fish kidney-derived reads, to remove sequences having high sequence identities to rainbow trout, the sequences were mapped locally using megablast^66^, to rainbow trout genome^67^ and transcriptome assemblies^28^. After removal of trout sequences (e value cutoff = 5e^−5^), assembly of high quality unmapped reads was performed using the CLC Bio Genomics Workbench 6.5.1 at an optimized k-mer size and assembled sequences clustered with TIGCL^68^ and CAP3^69^ with a minimal overlap of 100 bp and identity cut off of 95% to form unigenes. Sequence reads from the *T. bryosalmonae* spore sac library, were not filtered due to the absence of bryozoan genome / transcriptome assemblies and were directly assembled into contigs without removal of any contaminating bryozoan reads. All sequence reads have been deposited at the European Nucleotide Archive (ENA) under the accession number PRJEB19471. Fish kidney-derived contigs were further filtered by dc-megablast (e value cutoff = 5e^−3^) using two available myxozoan genome assemblies. Firstly, the current genome assembly of the myxosporean, *T. kitauei* (GCA_000827895.1) and, secondly, a partial assembly of the *T. bryosalmonae* genome (H. Hartikainen, B. Okamura, unpublished data). Contigs mapping to both genome assemblies constituted the final trout-derived transcriptome dataset. Owing to the low filtering capability of the *T. kitauei* genome, only the partial *T. bryosalmonae* genome assembly was used to filter the spore sac-derived contigs. Transcriptome metrics were calculated using Quast (version 3.1)^70^, the completeness of each transcriptome assembly was assessed using BUSCO (version 3)^71^ and raw reads were re-aligned to the assembly using Bowtie (version 1.1.1)^72^. Reciprocal blast (dc-megablast) analysis (e value cutoff = 5e^−3^) was undertaken to determine contigs in common with both transcriptome datasets (intersect contigs).

For assessment and visualisation of contamination, taxonomic assignment of each contig was carried out using BlobTools (version 0.9.19) at each stage of transcriptome production^73^.

### Transcriptome annotation

Annotation was performed on all contigs derived from *T. bryosalmonae* spore sacs and on myxozoan-filtered fish kidney-derived contigs using the Trinotate pipeline (version 2.0.2)^74^. Protein prediction was performed using TransDecoder (version r20140704) using default parameters^75^. Contigs were compared to Swiss-Prot and TREMBL (uniref90) databases^76–78^; downloaded March 2015 from www.uniprot.org) using NCBI BLAST+ (version 2.2.27) with *E*-values more than 10^−3^ considered non-significant. Both sets of contigs were also subjected to BLASTX searches against the non-redundant (nr) nucleotide database (downloaded from www.uniprot.org on the 17-03-2015). All contigs predicted to be protein coding were submitted to signal-P (version 4.1)^76^, TMHMM (version 2.0c)^77^ and to HMMER (version 3.0)^78^ against the PFAM database (version 27.0 downloaded Jan 2015)^79^. RNAmmer (version 1.2) was used to predict the presence of any contaminating ribosomal RNA sequences^80^. Gene Ontology assignment was performed using eggNOG, and SwissProt matches mapped to Gene Ontology classes using the mapping file: ftp://ftp.ebi.ac.uk/pub/databases/GO/goa/UNIPROT/gene_association.goa_uniprot.gz. GO classifications of intersect contigs were visualized using WEGO software^30^. Contig sequences were compared to the MEROPS (release 12.1; https://www.ebi.ac.uk/merops/download_list.shtml)^81^ database using BLASTX (version 2.2.31)^82^ with an e value cut off of 1e-05 and a percent identity cut off of 50%, with the most significant hit determined by the lowest e value.

### Real-time quantitative PCR (RT-qPCR)

We employed RT-qPCR to reveal parasite transcriptional differences between the two hosts. Total RNA representing uninfected / overtly infected bryozoans and uninfected/infected rainbow trout kidney tissue with a swelling grade of 2 (n= 3-4) was reverse transcribed into cDNA, as described previously^31^. Negative controls (without reverse transcriptase; -RT) were included to account for any residual DNA contamination.

Owing to limited amounts of bryozoan tissue, two batches of bryozoan and trout cDNA were utilised in this study providing 3 or 4 biological replicates per group. For bryozoan and trout cDNA, *ca* 1.5 µg and 5 µg of total RNA was reverse transcribed in 20 and 40 µl reactions respectively. Primers were designed to transcripts encoding proteins involved in; protein and lipid metabolism, developmental processes, cell-cell communication, cnidarian-specific structures or unknown / unannotated proteins (Supplementary Table S5). Primers were designed towards the 3’ end of each open reading frame aided by the considerable difference in codon usage between *T. bryosalmonae* and host genes (*ca*. 60-75% AT and 45-55% AT respectively). To facilitate primer design for some contigs, further sequence was obtained by anchored PCR and all primers were tested for specificity and efficiency using an in-house full-length spore sac cDNA library. Primer dimerization and melting profiles were assessed using OligoAnalyzer3.1 (https://www.idtdna.com/calc/analyzer). All primer pairs were confirmed to amplify only a single amplicon. Purified PCR amplicons, in TE buffer, were quantified using a NanoDrop 8000 Spectrophotometer, adjusted to 1 nM, and serially diluted as described previously^31^. Following sequence verification of each PCR standard, primer efficiency and concentration of each gene transcript was calculated by reference to its standard curve with efficiencies ranging from 1.80-1.96 (90-98%). Similarly, primers were designed for three selected reference genes to enable transcriptional normalization, namely; RPL18, RPL-12, and ELF-1α. RPL-12 has been shown to be a suitable reference gene for a range of invertebrate phyla, including cnidarians^83,84^. Likewise, RPL-18 and EF-1α have been shown to be highly consistent as reference genes in invertebrates^83,85^.

Relative quantification of *T. bryosalmonae* transcripts in each infected host was calculated using data from serially diluted reference DNA and normalized against the *T. bryosalmonae* reference gene, RPL18 in each PCR run and expressed as arbitrary units, as described previously^31^. Relative expression calculated in this way allows only comparison of host expression profiles for individual transcripts. Fold difference was used to facilitate interpretation of expression differences between hosts, with infected fish kidney expression levels normalized to infected bryozoan levels using the 2(-Delta Delta C(T)) method^86^.

In support of the validity and reliability of this RTq-PCR assay, our previous laboratory studies indicate that the unknown fish-specific secretory antigen, P14G8 is expressed at the protein level in fish kidneys from early (grade 0-1) to advanced (grade 2-3) clinical disease and closely correlates with P14G8 transcription in kidney samples (Holland et al., in preparation). In contrast, the much lower P14G8 expression in infected bryozoan samples was concomitant with a lack of detectable protein. The unknown secretory antigen C-39373, examined in this study, exhibited a similar fish-specific transcriptional profile as PG148 (Fig. 8C). The reliability of our RT-qPCR data is further supported by the apparent dominance of parasite LDL-R transcripts in myxozoan-infected fish tissues, as reported previously (Fig. 6 and Supplementary Table S4)^18,19^.

### DNA extraction and further verification of gene origin

Genomic DNA was extracted from Trizol-reagent homogenate using a DNA back extraction buffer (BEB) and dissolved in TE buffer, as described previously^87^. To further verify that the *T. bryosalmonae* transcripts corresponding to homeobox genes and a frizzled receptor homologue were parasite in origin, PCR analysis was conducted on genomic DNA from uninfected and *T. bryosalmonae*-infected bryozoan and rainbow trout kidney tissues using validated primers designed to specifically amplify genomic DNA, as listed in Supplementary Table S5. All PCR amplicons were sequenced verified, as described above.

### Statistical Analysis

All data sets were log2 transformed prior to statistical analysis. Using the Mann Whitney U test, *T. bryosalmonae* gene expression was considered significantly different between hosts when P <0.05. All statistical analyses were performed using GraphPad Prism (version 5.04, GraphPad Software Inc., La Jolla, USA).

## Author contributions

JWH took the lead in designing the study with input from CJS, MF, SS, EPA. JWH took the lead in writing of the manuscript with MF. JWH, ZQ, SY, and MF prepared tissues and RNA for library preps and RTqPCR analysis. JWH oversaw development of transcriptome assemblies and bioinformatic input by SS, EPA, and MF. MF, SS, and JWH compiled final data sets and figures/tables. JWH and CJS oversaw RTqPCR work and data analysis initially spearheaded by SY and later by MF, and BW. BO and HH provided bryozoan tissue samples for library preparation/expression studies and the partial parasite genome assembly for sequence filtering. All authors read, revised, and approved the final manuscript.

## Acknowledgements

This study was supported financially by the Biotechnology and Biological Sciences Research Council (BBSRC-1087-CS), a Swiss National Science Foundation Sinergia Project (CRS113 147649), a Ph.D. studentship to Marc Faber from the EU H2020 (H2020-SFS-10a-2014) program (ParaFishControl; 634429) and the Natural History Museum, London. We would like to thank Christopher Saunders-Davies of Test Valley Trout Ltd and Oliver Robinson of the British Trout Association for provision of fish and sampling facilities. Dr Daniel MacQueen of the Roslin Institute, University of Edinburgh, for provision of rainbow trout transcriptome assemblies. This publication reflects the views only of the authors, and the European Commission cannot be held responsible for any use which may be made of the information contained therein.

## Data availability

Additional data that support the findings of this study are available in; (1) Supplementary Tables S1-S5, (2) European Nucleotide Archive (ENA) sequence submission under the accession number PRJEB19471, (3) Figshare submission of intersect transcriptome (www.figshare.com, 10.6084/m9.figshare.11889672), (4) GenBank submission of individual contig sequences (accession numbers provided in Supplementary Table S5).

